# Plant trait heterosis is quantitatively associated with expression heterosis of the plastid ribosomal proteins

**DOI:** 10.1101/2021.02.16.431485

**Authors:** Devon Birdseye, Laura A. de Boer, Hua Bai, Peng Zhou, Zhouxin Shen, Eric A. Schmelz, Nathan M. Springer, Steven P. Briggs

## Abstract

The use of hybrids is widespread in agriculture, yet the molecular basis for hybrid vigor (heterosis) remains obscure. To identify molecular components that may contribute to the known higher photosynthetic capacity of maize hybrids, we analyzed proteomic and transcriptomic data from seedling leaf tissues of the hybrid, B73×Mo17, and its inbred parents. Subunits of complexes required for protein synthesis in the chloroplast and for the light reactions of photosynthesis were expressed above mid-parent and high-parent levels, respectively. Nuclear and plastid-encoded subunits were expressed similarly and in a dominant pattern with B73 as the high-parent for most proteins. The reciprocal hybrid displayed the same pattern with B73 still the dominant parent. Protein expression patterns were mostly the same in the adult leaf blade. To rank the relevance of expression differences to trait heterosis, we compared seedling leaf protein levels to adult plant heights of 15 hybrids. Expression heterosis (hybrid/mid-parent expression levels) was most positively correlated with plant height heterosis (hybrid/mid-parent plant height) for the plastid-encoded and nuclear-encoded chloroplast ribosomal proteins; the cytosolic ribosomal proteins were slightly less correlated. Ethylene biosynthetic enzymes were expressed below mid-parent levels in the hybrids, and the ethylene biosynthesis mutant, *acs2/acs6*, largely phenocopied the hybrid proteome, indicating that a reduction in ethylene biosynthesis may mediate the differences between inbreds and their hybrids. Levels of jasmonic acid biosynthetic enzymes were reduced in both *acs2/acs6* and the hybrid, and expression heterosis levels of these proteins were the most negatively correlated with plant height heterosis.

**Significance Statement:** Heterosis (hybrid vigor) boosts the productivity and resilience of crops and livestock above the levels of both parents, yet its underlying mechanisms remain unknown. We analyzed expression patterns of proteins in maize hybrids and their inbred parents. Differences in several molecular machines and biochemical pathways were found and quantitatively assessed using a panel of 15 hybrids. Seedling leaf chloroplast ribosomal proteins were able to quantitatively infer levels of adult plant heterosis. Expression levels of biosynthetic enzymes for the stress hormone, ethylene, were reduced in hybrids as was previously reported for the dicot Arabidopsis. Mutation of these genes in a maize inbred caused the proteome to resemble a hybrid. Repression of ethylene biosynthesis may be a conserved component of heterosis physiology.

## Main Text

### Introduction

Plants, animals, and humans display inbreeding depression associated with increased genetic uniformity and characterized by reduced vigor ^1^. Mating between genetically distinct inbred parents can produce hybrid vigor, or heterosis, defined as the difference in vigor between a hybrid and (a) the average of its parents or (b) the better-performing parent. Hybrids may out-perform their parents in terms of size, vigor, yield, abiotic and biotic stress resistance, longevity, and reproductive advantage ^2,3^. Maize is the most productive crop in the United States, and it was the first hybrid crop to be made and sold. Heterosis in maize increases many traits far above the levels of the higher parent, including biomass and harvestable grain ^4^. Despite its importance in agriculture, the changes in physiology that cause hybrid vigor remain obscure. Breeding hybrids requires expensive and labor-intensive field tests of both the hybrids and their parents. Progress in maize breeding has greatly improved yield through increased tolerance of high-density planting stress and this primarily constitutes the non-heterosis portion of yield ^5^. It is unclear whether substantial improvements in heterosis are possible and yet have lagged because breeders lack seedling biomarkers for adult plant heterosis.

QTL mapping has shown that minor effect loci for heterosis are distributed throughout the genome ^1^. Transcriptome ^6^ and proteome ^7^ profiles indicate that gene expression levels in hybrids are generally the average of their parents. A minority of mRNAs and proteins are expressed above or below mid-parent levels and most of these gene products are not obviously related to each other in function or to hybrid phenotypes, making their unusual levels difficult to interpret ^8^. Despite these ambiguities, progress has been made in understanding hybrid vigor.

Recent work has implicated the plant hormone ethylene (ET) in heterosis. In Arabidopsis hybrids, ET is produced at reduced levels and application of exogenous ET reduced hybrid vigor ^9^. Thus, ET potentially mediates at least some aspects of heterosis, though it is unclear which proteins are affected by reduced ET biosynthesis.

Maize hybrids have greater photosynthetic capacity than their inbred parents ^10^, which presumably contributes to their increased biomass and yield. In both maize and Arabidopsis, various genes for photosynthesis have been observed to be expressed above mid-parent levels ^10–12^. Some metabolites and enzymes of the photosynthetic carbon reactions are elevated in maize hybrids while others in photorespiration are repressed ^13^. We sought to identify protein biomarkers in seedling leaves that contribute new insights to the physiology of hybrid vigor and can be used to predict levels of adult trait heterosis.

## Results

### Dominant and Overdominant Expression Patterns of Chloroplast Protein Complexes

Multiplexed proteomic analyses were performed using TMT peptide tags and high-resolution mass spectrometry to quantify differences in the levels of proteins extracted from seedling leaves of hybrids and their inbred parents. Expression levels in the hybrid, B73×Mo17, were compared to high-parent (HP) levels, low-parent (LP) levels, and calculated mid-parent (MP) levels to identify non-additive expression. Readers can use our website to visualize expression of chosen genes in the hybrid and inbred parents (https://devonbirdseye.shinyapps.io/ExpressionViewer/).

Of 10,141 measured leaf proteins, 638 (6%) were expressed above HP. Functional enrichment performed using The Database for Annotation, Visualization and Integrated Discovery (DAVID) ^14,15^ revealed that these were enriched for chloroplast-localization. Of the 1,068 chloroplast-localized proteins observed, 256 (24%) were expressed above HP levels. This included 65 of the Photosynthesis-Associated Nuclear Genes (PhANGs) and Photosynthesis-Associated Plastid Genes (PhAPGs). The PhAPG/PhANG proteins comprise photosystem I, photosystem II, cytochrome b6/f complex, NAD(P)H dehydrogenase-like complex, ATP synthase, and the carbon-fixation enzyme, Rubisco. PhANG and PhAPG proteins were expressed 19% above HP levels, on average (Figure 1A). The remaining 189 chloroplast-localized proteins were weakly enriched for carbon fixation.

**Figure 1.**
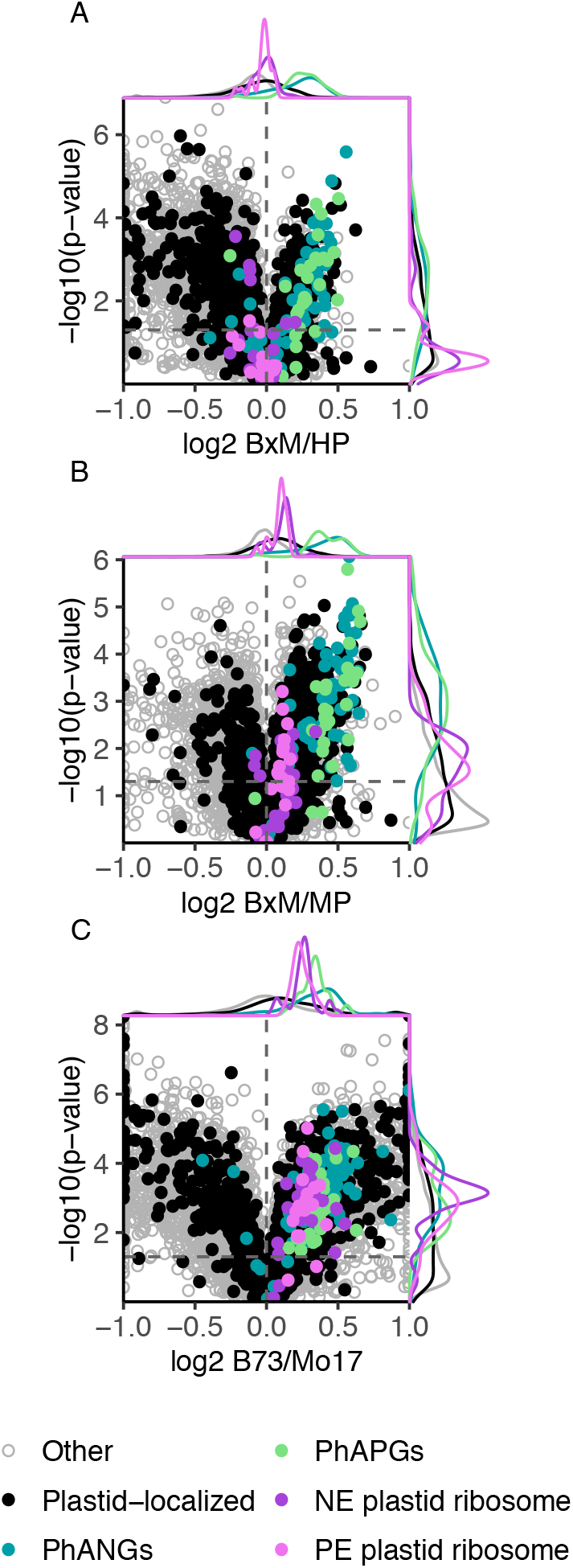
Volcano plots displaying expression patterns of the most significantly non-additive proteins, representing B73×Mo17/high-parent (A), B73×Mo17 /mid-parent (B), and B73/Mo17 (C). Photosynthesis-associated Nuclear Genes (PhANGs), Photosynthesis-associated Plastid Genes (PhAPGs), Nuclear-Encoded (NE) plastid ribosome, and Plastid-Encoded (PE) plastid ribosome proteins are color-coded.

Expression between MP and HP levels was observed for 956 proteins (9%). They were enriched for plastid ribosomal proteins, encoded by either the nuclear or plastid genome. The ribosome subunits were expressed 8% above MP, on average (Figure 1B). Less enrichment was observed for enzymes of carbon metabolism. While not enriched, carbon fixation-related proteins were differentially expressed in the hybrid, some above MP and some below (Figure S1).

Proteins that were expressed non-additively were frequently expressed at different levels between the parents, and expression in B73 was generally higher than in Mo17 (Figure 1C). However, some proteins such as photosynthetic electron transfer C, ribose-5-phosphate isomerase, and the oxygen evolving complex assembly proteins were expressed higher in Mo17 (Dataset S1).

Expression below LP levels was observed for 517 proteins (5%). They were weakly enriched for amino acid biosynthesis, fatty acid degradation, chaperone proteins, and peroxisomal proteins. Expression between MP and LP was observed for 1,222 proteins (12%). These showed weak enrichment for chaperone proteins and protein processing in the ER.

The non-additive patterns of expression seen in the proteome of juvenile leaves were also observed in the blades of mature leaves (Figure S2 A). However, there was a reversal of relative expression levels between the parents for plastid ribosomal proteins (Figure S2 B). B73 was high parent in the seedling leaf, whereas Mo17 was high parent in the leaf blade.

### Correlations between Trait Heterosis and Expression Heterosis

To rank the relationships between trait heterosis and protein expression heterosis, we examined six hybrids generated from diverse inbreds. B73×Mo17 and its reciprocal hybrid, Mo17×B73 (the female parent is listed first), displayed the same differential expression patterns, indicating that the patterns are caused by combining the nuclear genomes, with little or no effect due to maternal inheritance of the plastid genome (Figure 2 C–D). Inbreds B73 and B84 are related members of the Stiff Stalk pool of germplasm. Inbreds Mo17 and A682 are part of the Non-Stiff Stalk pool. Hybrids made from crosses between these pools have strong heterosis in contrast to hybrids made from crosses within each pool ^16^. The highest expression heterosis of the PhANG/PhAPG proteins was observed in hybrids with the least heterosis (Figure 2 E–F). In contrast, the plastid ribosomal proteins were expressed at or below HP levels in the hybrids with low levels of heterosis and substantially above HP levels in the hybrid with the greatest heterosis (Figure 2).

**Figure 2.**
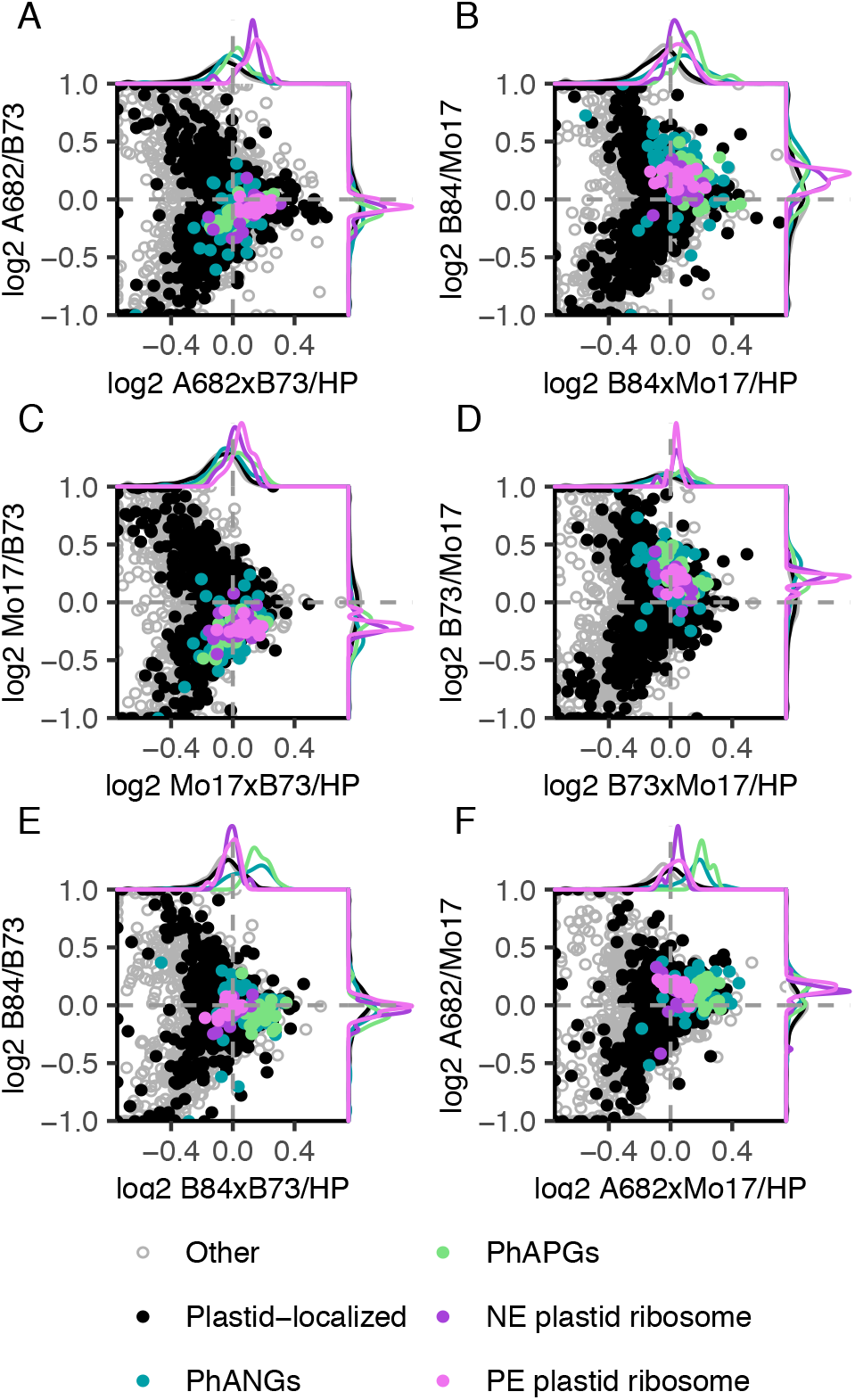
Volcano plots displaying plastid-localized proteins in seedling leaves of six hybrids relative to high-parent (HP) levels. Hybrids represented in A-F are ordered from greatest plant height heterosis (A) to least plant height heterosis (F). Each data point represents the mean of three biological replicates. Photosynthesis-associated Nuclear Genes (PhANGs), Photosynthesis-associated Plastid Genes (PhAPGs), Plastid-Encoded (PE) plastid ribosomes, and Nuclear-Encoded (NE) plastid ribosomes are color-coded.

We additionally analyzed nine hybrids made by back-crossing four IBM recombinant inbred lines (RILs) to both parents, B73 and Mo17. The RIL hybrids were approximately 50% as heterozygous as the parent hybrid and were selected based on their varied levels of plant height heterosis (Dataset S2 and Dataset S3). We examined collectively these 15 hybrids and their parents, including the RIL hybrid set and the six hybrids described above. As a measure of trait heterosis, we utilized plant height, which is correlated with grain yield ^17^. We calculated the Pearson’s correlations between expression heterosis and plant height heterosis (hybrid height/MP height). Readers can obtain Pearson correlation values for genes of interest using our website (https://devonbirdseye.shinyapps.io/ExpressionViewer/). It is important to note that there was no correlation between expression levels and plant height *per se*; only the hybrid/mid-parent values for expression and height were correlated. This underscores the specificity of our biomarkers for the trait of heterosis.

The individual Pearson correlations between expression heterosis levels of 581 proteins and plant height heterosis was greater than 0.5 (Dataset S4). This set of proteins was enriched for the terms ribosome (93 proteins), chloroplast (48 proteins), protein biosynthesis (20 proteins), photosynthesis (11 proteins), plastid chromosome (5 proteins), and tetratricopeptide repeat (12 proteins) (Figure 3). Of the top 1.5% of proteins whose expression heterosis was most correlated with plant height heterosis, 18 out of 54 were from the plastid ribosome. Pearson’s correlations of 0.79-0.91 were observed for these proteins, half of which were nuclear-encoded and half were plastid-encoded. Correlations were most positive for the plastid ribosomal proteins, slightly less positive for the cytosolic ribosomal proteins, and moderately positive for the PhANG/PhAPG proteins (Figure 3). There were 413 proteins with a Pearson’s correlation less than −0.5; these were most strongly enriched for oxidoreductase (39 proteins), biosynthesis of secondary metabolites (63 proteins), protease (18 proteins), biosynthesis of antibiotics (32 proteins), and alpha-linoleic acid metabolism (7 proteins) terms (Figure 3).

**Figure 3.**
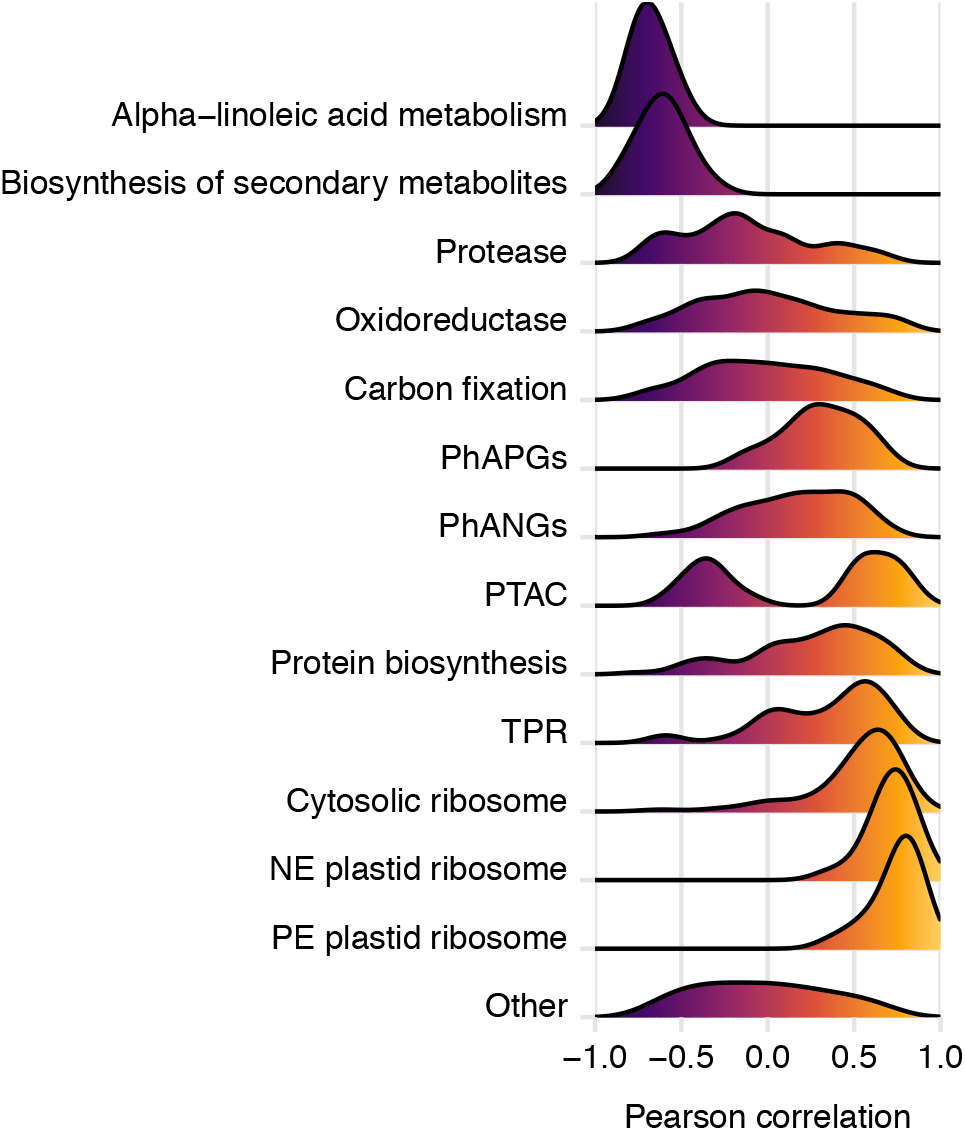
Figure 3. Density curves of Pearson correlations between protein expression heterosis and plant height heterosis in the combined RIL and six hybrids datasets.

### Discordance between Transcript and Protein Expression Heterosis

The proteome differences between chloroplasts of the hybrid and its inbred parents were not mirrored in their nuclear transcriptomes (Figure S3). The nuclear transcripts for PhANGs were between MP and HP. Nuclear transcripts for plastid ribosomal subunits were substantially below MP in contrast to their proteins which were above MP. Most PhANG transcripts were expressed higher in B73, consistent with their protein expression. However, most plastid ribosomal subunit transcripts were higher in Mo17, strikingly discordant with parental expression of their proteins.

Expression of nuclear transcripts for the plastid ribosome relative to MP levels was less strongly correlated with plant height heterosis than that of their proteins, and expression relative to HP levels was even weaker (Figure S4). PhANG transcript expression relative to MP levels was substantially more positively correlated than that of their proteins, while expression relative to HP levels had a weakly positive correlation, similar to their proteins. Transcripts for the cytosolic ribosome had no correlation with plant height heterosis, unlike their proteins. Plastid-encoded transcripts were not measured.

Overall, 640 transcripts were expressed above HP, with functional enrichment for circadian rhythm accounting for 7 transcripts; the remaining transcripts had no functional enrichment. With a few exceptions, most circadian-related transcripts were expressed above MP levels (Figure S5A), while expression heterosis of these transcripts varied in their correlations with plant height heterosis (Figure S5B). For example, phytochrome B (PhyB) transcripts were positively correlated while PHYTOCHROME-INTERACTING FACTOR 3 (PIF3) transcripts were negatively correlated. There were 40 transcription factors expressed above HP, including 5 ethylene-responsive factors, 4 HOX-domain proteins, 1 auxin-responsive factor, and the circadian regulator, LHY. Conversely, 517 transcripts were expressed below LP, with functional enrichment for metabolic pathways, especially biosynthesis of amino acids. Only one transcription factor, PHD16, was expressed below LP.

### Expression Heterosis is Phenocopied by an *acs* Mutant

Mutations in two of the maize genes encoding ACC synthase have been functionally characterized (*Zmacs2* and *Zmacs6*) and were shown to have decreased endogenous ET levels ^18–21^. Transcripts for ZmACS2, ACO, and SAM1/2 homologs, were found to be expressed substantially below MP levels (Figure S6), whereas the ZmACS6 transcript was not detected. Several isoforms of enzymes predicted to be involved in ET biosynthesis were also expressed significantly below MP levels, although it is unclear if reduced expression of these enzymes would cause a reduction in ET levels.

Proteomic analysis of the ET biosynthesis double mutant, *Zmacs2/6*, in the B73 genetic background ^18^ revealed that the mutant phenocopied the hybrid molecular phenotype. Proteins expressed above or below MP in A682×B73 (the hybrid with the greatest heterosis), were expressed similarly in the mutant relative to the inbred, B73 (Figure 4A). Protein levels of most PhANGs/PhAPGs and plastid ribosome subunits were elevated in the mutant at equal proportions as in A682×B73 (Figure 4B). The exceptions were two ATP synthase subunits and two plastid ribosomal proteins, which were significantly above MP in the hybrid but unchanged in the mutant. Expression levels of jasmonic acid (JA) biosynthesis enzymes were reduced in the *Zmacs2/6* mutant relative to B73 and in hybrids relative to MP (Figure S7). Of the 538 proteins expressed above MP in A682xB73, only 18 were repressed in *Zmacs2/6*; these were enriched for biosynthesis of secondary metabolites and also included three PLASTID TRANSCRIPTIONALLY ACTIVE CHROMSOME (PTAC) proteins. Of the 285 proteins expressed below MP in A682xB73, only 15 were elevated in *Zmacs2/6*; no functional enrichment was found for these.

**Figure 4.**
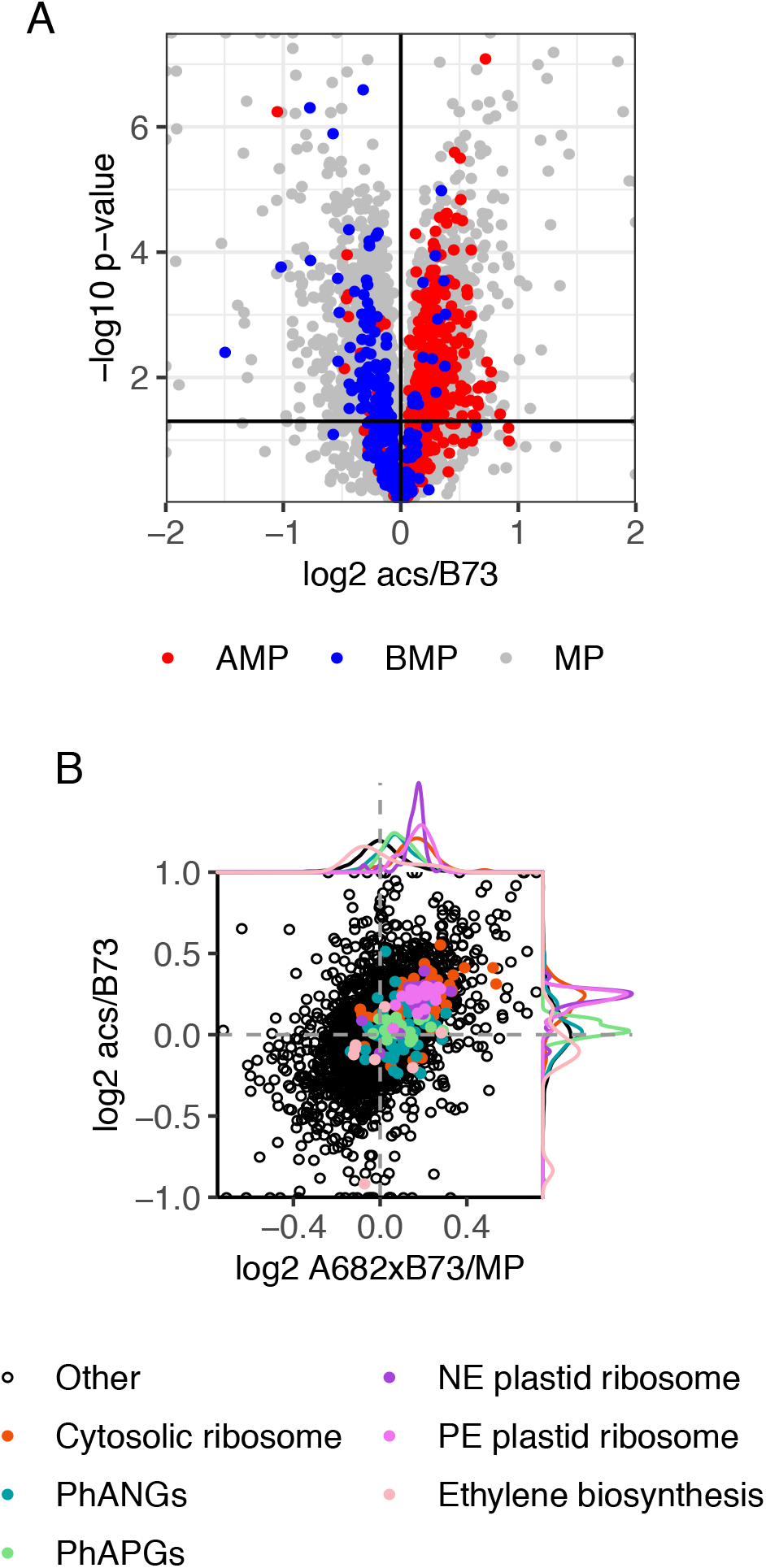
(A) Volcano plot of protein expression in the *acs2-6* double mutant in B73 background relative to B73. Each data point represents the mean of five biological replicates. Colors correspond to their expression in the hybrid with greatest plant height heterosis, A682×B73, designated as mid-parent (MP), above mid-parent (AMP), or below mid-parent (BMP) (B) Plastid-localized proteins in seedling leaves of the *acs2-6* double mutant in B73 background relative to B73, compared to expression in A682×B73 relative to MP. Each data point represents the mean of five biological replicates for *acs2-6*/B73 and three biological replicates for the hybrid and its inbred parents. Photosynthesis-Associated Nuclear Genes (PhANGs), Photosynthesis-Associated Plastid Genes (PhAPGs), Plastid-Encoded (PE) plastid ribosomes, Nuclear-Encoded (NE) plastid ribosomes, and ethylene biosynthesis proteins are color-coded.

## Discussion

Most plastid-localized proteins were expressed at mid-parent levels in the hybrid, B73×Mo17. Non-additive expression patterns were observed for a minority of proteins and RNAs. Photosynthesis-related proteins and plastid ribosomal proteins represented the major groups expressed above HP and MP levels, respectively. For both groups, the nuclear-encoded and plastid-encoded subunits were expressed similarly. Elevated expression of photosynthetic proteins in hybrids is diurnally-regulated, with highest levels in the morning ^13^, which corresponds to the sampling time in our experiments. Many transcription factors were also expressed above HP. Proteins and transcripts expressed below LP were enriched most strongly for amino acid biosynthesis, consistent with previous measurements of hybrid metabolite levels ^13^. We observed that circadian-related genes were enriched in the transcripts expressed above HP, consistent with previous reports of an altered circadian rhythm in hybrids ^9,10,22^.

We hypothesize that coupling of photosynthesis to growth - specifically, the heterosis portions of each - may be through the plastid ribosome, which was positively correlated with plant height heterosis at both protein and transcript levels. The cytoplasmic ribosome is nearly as well correlated and is also likely to contribute to heterosis traits. Subunits of the cytoplasmic ribosome are expressed above MP in the maize seminal root ^23^. In yeast, proteome reallocation to ribosomes enables faster growth in rich media ^24^. Elevated photosynthetic capacity may be important for heterosis even though it does not appear to be quantitatively coupled to the levels of heterosis. Hybrids made from inbreds of the same heterotic pool expressed the PhANG and PhAPG proteins above high-parent levels whereas PhANG transcripts were higher in hybrids made from crosses between heterotic pools. Consequently, PhANG transcripts were quantitatively associated with plant height heterosis, while their protein levels were not. Nevertheless, hybrids have greater photosynthetic capacity than their parents and this may be necessary to produce the greater biomass and yield of hybrids ^10^.

Other groups of proteins with positive correlations to heterosis included PTAC proteins and protein biosynthesis-related proteins, consisting of elongation factors and initiation factors. Elevated expression of both of these groups of proteins in the hybrid may contribute to the hybrid-specific expression patterns of the chloroplast- and nuclear-encoded proteins. Tetratricopeptide repeat (TPR) proteins were enriched in the positively correlated group, and some of these are known to mediate protein-protein interactions and play roles in stress and hormone signaling ^25^.

Groups of proteins enriched amongst those with negative correlations included oxidoreductase, consisting of several dehydrogenases, lipoxygenases, and one of the ethylene biosynthetic enzymes, ACC oxidase. Proteases were negatively correlated. The two most negatively correlated groups were biosynthesis of secondary metabolites and alpha-linoleic acid metabolism, consistent with the proposed tradeoff between stress responses and plant growth. Carbon fixation-related proteins had a mixed pattern of expression differences in the hybrid, and overall were neither positively nor negatively correlated with plant height heterosis. Others report that hybrids are associated with either elevated levels of photosynthetic proteins (hybrid ZD909) or stress proteins (hybrid ZD808) but not both ^26^.

Our results indicate that the physiology of heterosis may be conserved between monocots and dicots. ET biosynthetic enzyme gene expression was repressed in maize hybrids as was previously reported for Arabidopsis hybrids ^9^. Reduced expression of ET biosynthetic enzymes caused by the *Zmacs2/6* mutation in B73, phenocopied the heterosis expression levels of chloroplast proteins. Traits of the maize *acs2* and *acs6* mutants are known to resemble those of hybrids; these include transpiration, stomatal closure, carbon dioxide assimilation, and elevated levels of foliar chlorophyll, Rubisco, and soluble protein ^18^. JA biosynthetic enzyme levels were repressed in both the hybrid and in the *Zmacs2-6* mutant, indicating that JA biosynthesis may be regulated by ET. Interestingly, the JA biosynthetic pathway was enriched in the proteins that were negatively correlated with plant height heterosis, suggesting a potential role for reduced JA biosynthesis. The most notable differences between A682×B73 (the hybrid with the greatest heterosis) and the *Zmacs2-6* mutant were expression levels of three PTAC proteins which were elevated in the hybrid and repressed in the mutant. Levels of these plastid RNA polymerase protein subunits were positively correlated with plant height heterosis.

The discrepancy between accumulation patterns of transcripts and their proteins argues that post-transcriptional mechanisms are selectively regulating protein levels in the inbreds and their hybrids. Developmental and circadian patterns of expression can uncouple transcripts from their proteins ^27^. Expression of plastid ribosomal proteins was particularly interesting. They were expressed above MP in B73xMo17 while their transcripts were below MP. Across a panel of 15 hybrids, expression heterosis of these proteins was highly correlated with plant height heterosis. Most nuclear-encoded and plastid-encoded subunits were expressed higher in B73 than in Mo17. In contrast, transcripts for the nuclear-encoded subunits were mostly expressed higher in Mo17. It’s unclear how B73 accumulates more of each ribosomal protein from substantially lower transcript levels than in Mo17. The levels of PhANG and PhAPG proteins were higher in B73 than in Mo17, in parallel with their nuclear transcript levels. Thus, for the digenomic protein complexes of the chloroplast, one parent is dominant for expression of nuclear and plastid-encoded subunits. Their levels are higher in B73 and above MP or HP in the hybrid, even when their transcripts follow an opposite pattern. This is true whether B73 is the female or male parent. Proteome dominance has been observed in the non-photosynthetic immature ear of the maize hybrid, ZD909 ^7^. The authors found that many enzymes involved in carbon and nitrogen assimilation were expressed at high-parent levels. In Arabidopsis, transcripts for cell division are more highly expressed in one parent, and in the other parent transcripts for photosynthesis are higher. The hybrid expressed both sets above MP levels and the combination of the two pathways may be important for heterosis ^28^. Further work is needed to understand the mechanisms of expression dominance and the roles it may play in heterosis.

## Materials and Methods

### Plant growth and sampling

Plant materials were all grown in growth chambers for collection of seedling leaf tissues and grown in the field for collection of the mature leaf blade tissues. At least three biological replicates were used for each genotype. Details on the plant materials and sampling and phenotyping are described in SI Appendix, SI Materials and Methods.

### RNA-Seq analysis

Sequence libraries were prepared using the standard TruSeq Stranded mRNA library protocol and sequenced on NovaSeq 150bp paired end S4 flow cell to produce at least 20 million reads for each sample. CPM values were calculated (Dataset S6 and Dataset S7) and used to calculate averages and ratios. Details and procedures for RNA-Seq and data analysis are described in SI Appendix, SI Materials and Methods.

### Proteomics data acquisition and analysis

Spectra are acquired on a Q-exactive-HF mass spectrometer (Thermo Electron Corporation, San Jose, CA). Tandem-mass tag (TMT) abundances were normalized to the arithmetic mean of the B73 replicates in each run (Dataset S5) and used to calculate averages and ratios. Protein group assignment was based off of KEGG ^29^, CornCyc ^30^, and NCBI^31^ (Dataset S8). Details and procedures for proteomics methods and data analysis are described in SI Appendix, SI Materials and Methods.

## Acknowledgments

This material is based upon work supported by the National Science Foundation Grant No. 1546899 to S.P.B and N.M.S., and the National Science Foundation Graduate Research Fellowship under Grant No. DGE-1650112 to D.B. Any opinions, findings, and conclusions or recommendations expressed in this material are those of the authors and do not necessarily reflect the views of the National Science Foundation.

