## Supplementary material for "Plant trait heterosis is quantitatively associated with expression heterosis of the plastid ribosomal proteins": SI Appendix

*Steven P. Briggs

**This PDF file includes:**

Supplementary Materials and Methods
Figures S1 to S7
Legends for Datasets S1 to S8
SI References

**Other supplementary materials for this manuscript include the following:**

Datasets S1 to S8

### Supplementary Materials and Methods

**Plant growth and sampling.** Plant materials were all grown in the green house for collection of seedling tissues and grown in the field for collection of adult tissues. The seedling leaf and mature leaf blade tissues of B73, Mo17 and B73xMo17 were collected in May of 2016 in St Paul, Minnesota, while tissues from the six hybrids and RIL hybrids experiments were collected from July to November of 2019 in St Paul, Minnesota. For seedling tissues, seeds were imbibed for 24 hours in distilled water and grown in growth chambers at 25℃ under 12h light-dark cycles to V2/V3 stage (day 9) for collection of the V2 leaves. Tissue from at least four plants was pooled for each replicate and the three biological replicates were sampled at the same time (12PM) on different dates in order to sample the same growth stage for each genotype. For the *acs2/6* mutant and B73 seedling leaf samples, seeds were imbibed for 48 hours and manually examined to select uniformly germinating seedlings based on radical emergence and the lack of mold. They were then grown in growth chambers at 25°C under 12h light-dark cycles to V2/V3 stage (day 10). The V3 leaf was collected for 5 biological replicates of each genotype.

**RNA-Seq analysis.** Sequence libraries were prepared using the standard TruSeq Stranded mRNA library protocol and sequenced on NovaSeq 150bp paired end S4 flow cell to produce at least 20 million reads for each sample. Both library construction and sequencing were done in the University of Minnesota Genomics Center. Sequencing reads were then processed through the nf-core RNA-Seq pipeline [1] for initial QC and raw read counting. In short, reads were trimmed by Trim Galore! (https://github.com/FelixKrueger/TrimGalore) and aligned to the B73 maize reference genome (Zm-B73-REFERENCE-GRAMENE-4.0) [2] using the variant-aware aligner Hisat2 [3] which also takes 90 million common variants to account for mapping bias. Uniquely aligned reads were then counted per feature by featureCounts [4]. Raw read counts were then normalized by library size and corrected for library composition bias using the TMM normalization approach [5] to give CPMs (Counts Per Million reads) for each gene in each sample, allowing direct comparison across samples (Supplemental Tables 6 and 7). Hierarchical clustering, principal component analysis and t-SNE clustering were used to explore sample cluster patterns and to remove or substitute bad replicates.

**Proteomics Sample Preparation.** Ground tissue powders were suspended in extraction buffer (8M Urea/100mM Tris/5mM Tris(2- carboxyethyl)phosphine (TCEP)/phosphatase inhibitors, pH 7). Proteins were precipitated by adding 4 volumes of cold acetone and incubated at 4oC for 2 hours. Samples were centrifuged at 4,000xg, 4oC for 5 minutes. Supernatant was removed and discarded. Proteins were re-suspended in Urea extraction buffer (8M Urea/100mM Tris/5mM TCEP/phosphatase inhibitors, pH 7) and precipitated by cold acetone one more time. Protein pellets were washed by cold methanol to further remove non-protein contaminants. Protein pellets were suspended in extraction buffer (8M Urea/100mM Tris/5mM TCEP/phosphatase inhibitors, pH 7). Proteins were first digested with Lys-C (Wako Chemicals, 125-05061) at 37oC for 15 minutes. Protein solution was diluted 8 times to 1M urea with 100mM Tris and digested with trypsin (Roche, 03 708 969 001) for 4 hours. Digested peptides were purified on a Waters Sep-Pak C18 cartridges, eluted with 60% acetonitrile. TMT-10 labeling was performed in 50% acetonitrile/150mM Tris, pH7. TMT labeling efficiency was checked by LC-MS/MS to be greater than 99%. Labeled peptides from different samples were pooled together for 2D-nanoLC-MS/MS analysis. An Agilent 1100 HPLC system was used to deliver a flow rate of 600 nL min-1 to a custom 3-phase capillary chromatography column through a splitter. Column phases were a 20 cm long reverse phase (RP1, 5 μm Zorbax SB-C18, Agilent), 6 cm long strong cation exchange (SCX, 3 μm PolySulfoethyl, PolyLC), and 20 cm long reverse phase 2 (RP2, 3.5 μm BEH C18, Waters), with the electrospray tip of the fused silica tubing pulled to a sharp tip (inner diameter <1 μm). Peptide mixtures were loaded onto RP1, and the 3 sections were joined and mounted on a custom electrospray adapter for on-line nested elutions. Peptides were eluted from RP1 section to SCX section using a 0 to 80% acetonitrile gradient for 60 minutes, and then were fractionated by the SCX column section using a series of 20 step salt gradients of ammonium acetate over 20 min, followed by high-resolution reverse phase separation on the RP2 section of the column using an acetonitrile gradient of 0 to 80% for 150 minutes.

**Proteomics data acquisition.** Spectra are acquired on a Q-exactive-HF mass spectrometer (Thermo Electron Corporation, San Jose, CA) operated in positive ion mode with a source temperature of 275 °C and spray voltage of 3kV. Automated data-dependent acquisition was employed of the top 20 ions with an isolation windows of 1.0 Da and collision energy of 30. The mass resolution is set at 60,000 for MS and 30,000 for MS/MS scans, respectively. Dynamic exclusion is used to improve the duty cycle.

**Proteomics data analysis.** The raw data was extracted and searched using Spectrum Mill vB.06 (Agilent Technologies). MS/MS spectra with a sequence tag length of 1 or less were considered to be poor spectra and were discarded. The remaining high quality MS/MS spectra were searched against a maize protein database including maize B73 (Zm-B73-REFERENCE-GRAMENE-4.0) and Mo17 (Zm-Mo17-REFERENCE-CAU-1.0) reference genomes [2,6]. A 1:1 concatenated forward-reverse database was constructed to calculate the false discovery rate (FDR). There were 435,212 protein sequences in the final protein database. Search parameters were set to Spectrum Mill's default settings with the enzyme parameter limited to full tryptic peptides with a maximum mis-cleavage of 1. Cutoff scores were dynamically assigned to each dataset to obtain the false discovery rates (FDR) of 0.1% for peptides, and 1% for proteins. Proteins that share common peptides were grouped using principles of parsimony to address protein database redundancy. Total TMT-10 reporter intensities were used for relative protein quantitation. Peptides shared among different protein groups were removed before TMT quantitation. Isotope impurities of TMT-10 reagents were corrected using correction factors provided by the manufacturer (Thermo). Median normalization was performed to normalize the protein TMT-10 reporter intensities in which the log ratios between different TMT-10 tags were adjusted globally such that the median log ratio was zero.

**Proteomics data deposition.** The raw spectra for the proteome data have been deposited in the Mass Spectrometry Interactive Virtual Environment (MassIVE) repository (massive.ucsd.edu/ProteoSAFe/static/massive.jsp) (accession ID MSV000085916). FTP download link before publication: ftp://MSV000085916@massive.ucsd.edu; FTP download link after publication: ftp://massive.ucsd.edu/MSV000085916/.

**Expression analysis.** For protein analyses, tandem-mass tag (TMT) abundances were normalized to the arithmetic mean of the B73 replicates in each run (Supplemental Dataset 5). For RNA analysis of the original seedling leaf and leaf blade, previously published counts per million (CPM) values were used [7] For six hybrid and RIL hybrid RNA analyses, CPM values were used (Supplemental Datasets 6 and 7 for six hybrids and RIL hybrids, respectively). For each experiment, genes or proteins were removed if they were not detected in all biological replicates. The arithmetic mean of the biological replicates was then calculated and used to calculate hybrid/mid-parent and parent/parent ratios. The log base 2 was then calculated for each ratio. A T-test was used to calculate p-values. Data was manipulated using the R packages dplyr [8], plyr [9], tidyr [10], reshape2 [11], stringr [12], and data.table [13]. Functional enrichment performed using The Database for Annotation, Visualization and Integrated Discovery (DAVID) [14,15]. Scatterplots and volcano plots were generated using ggplot2 [16], density plots were generated using cowplot [17], and plots were assembled into paneled figures using patchwork [18]. KEGG maps were generated using pathview [19], with the default settings in which the sum is calculated for each enzyme for expression maps and the absolute maximum calculated for the Pearson correlation map. Colors for CCA1 on the circadian rhythm maps were added manually. For correlation analyses, the Pearson correlations (Supplemental Dataset 4) were calculated from the hybrid/mid-parent expression ratios (Supplemental Dataset 1) to the hybrid/mid-parent plant height ratios (Supplemental Dataset 3) and plotted using ggridges [20]. All source codes used for expression analyses are available on Github: https://github.com/devonbirdseye/HeterosisManuscript

**Protein group assignment.** The plastid proteome was defined according to a previous report [21]. PhANGs were defined as the proteins in known chloroplast complexes using KEGG maps zma00195, excluding the electron transport proteins, zma00196, and CornCyc reactions 1.97.1.12 and 1.10.3.9 [22,23], all of which were downloaded on February 26, 2020. The PhAPGs were defined as the plastid-encoded proteins with NCBI annotations [24] as PSI, PSII, cytochrome b6f, and ATP synthase. Nuclear-encoded plastid ribosomal proteins were defined as those containing “30S” or “50S” in their NCBI annotation [24] as well as “plastid” or “chloroplast” in their maize-GAMER GO annotations [25], with term descriptions retrieved using the R package AnnotationDbi [26]. Cytosolic ribosomal proteins were defined as those containing “40S” or “60S” in their NCBI annotation, except for those containing “mitochondrial” or “biogenesis”. Ethylene biosynthetic proteins were defined using CornCyc reactions 2.5.1.6, 4.4.1.14 and 1.14.17.4 [23], downloaded on May 5, 2020. All accession conversions to v4 were made using the MaizeGDB accession conversion table, downloaded on May 18, 2020 [23]. Proteins involved in carbon fixation, biosynthesis of secondary metabolites, and alpha-linoleic acid metabolism were defined using KEGG maps zma00710, zma01110, and zma00592, respectively. TPR and PTAC proteins were defined as those containing “TPR” and “plastid transcriptionally active” in their NCBI annotation [24], respectively. Oxidoreductase and protease proteins were defined as those with the term GO:0016491 and GO:0008233 in their maize-GAMER GO annotations [25], respectively. Protein biosynthesis proteins were defined as those containing “elongation factor” or “initiation factor” in their NCBI annotation [24]. A complete list of gene group assignments and their sources is provided in Supplemental Dataset 8.

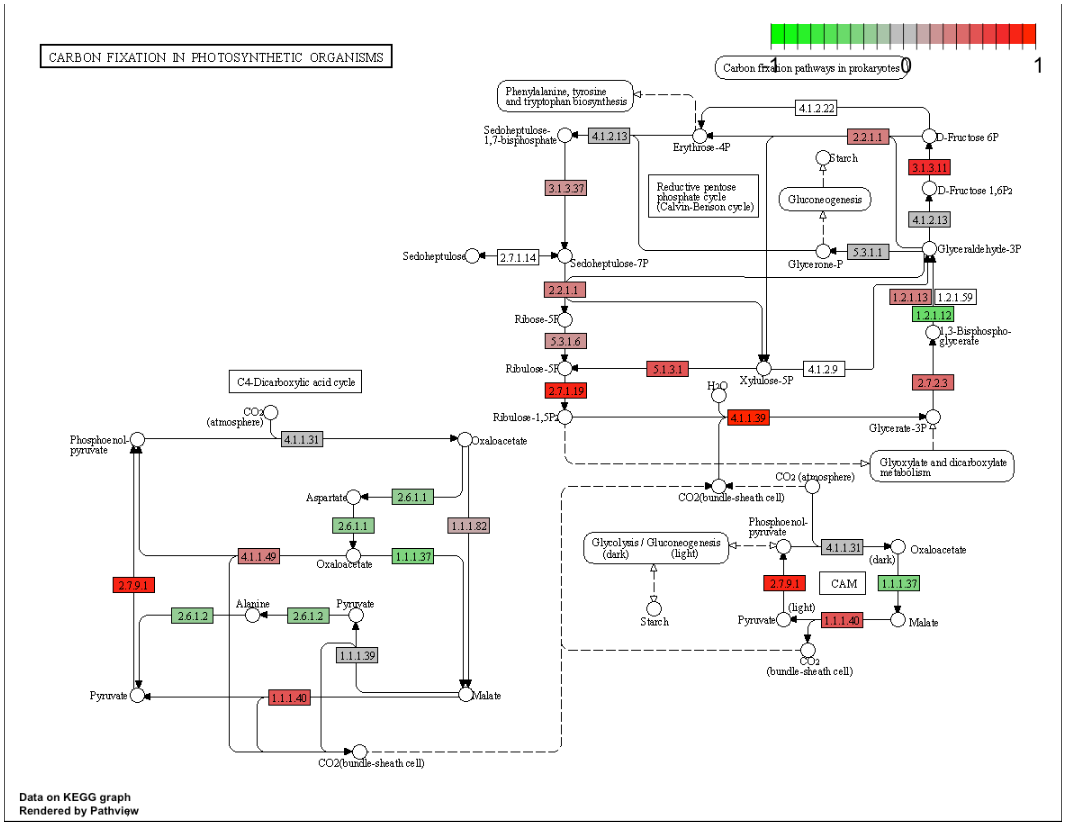


**Fig. S1.** Expression of carbon fixation enzymes in the hybrid B73xMo17 relative to its mid-parent level.

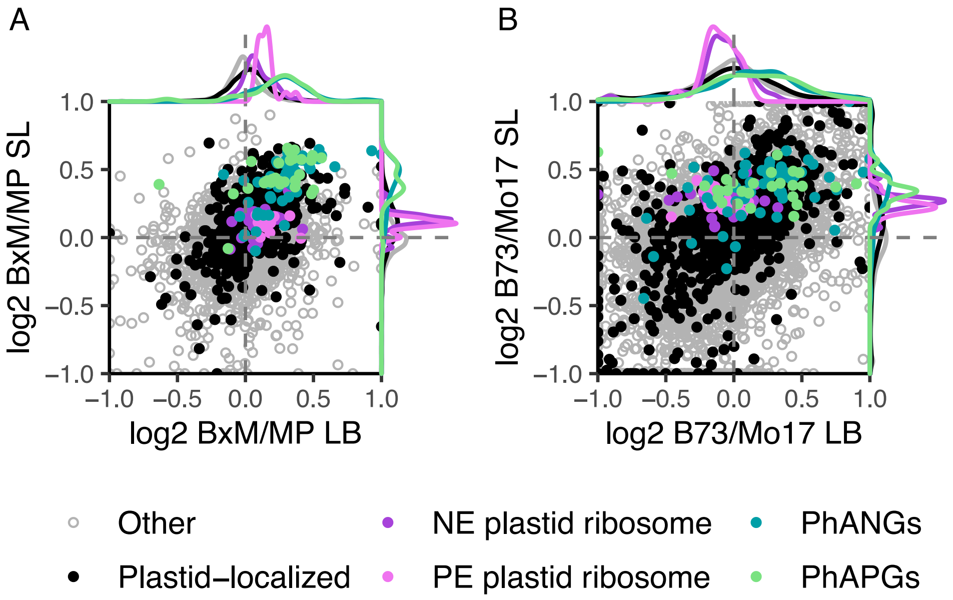


**Fig. S2.** Comparison of expression of the significantly non-additive proteins in seedling leaf versus leaf blade tissue. A compares hybrid/mid-parent values; B compares parent/parent values. Photosynthesis-associated Nuclear Genes (PhANGs), Photosynthesis-associated Plastid Genes (PhAPGs), Plastid-Encoded (PE) plastid ribosomes, and Nuclear-Encoded (NE) plastid ribosomes are color-coded.

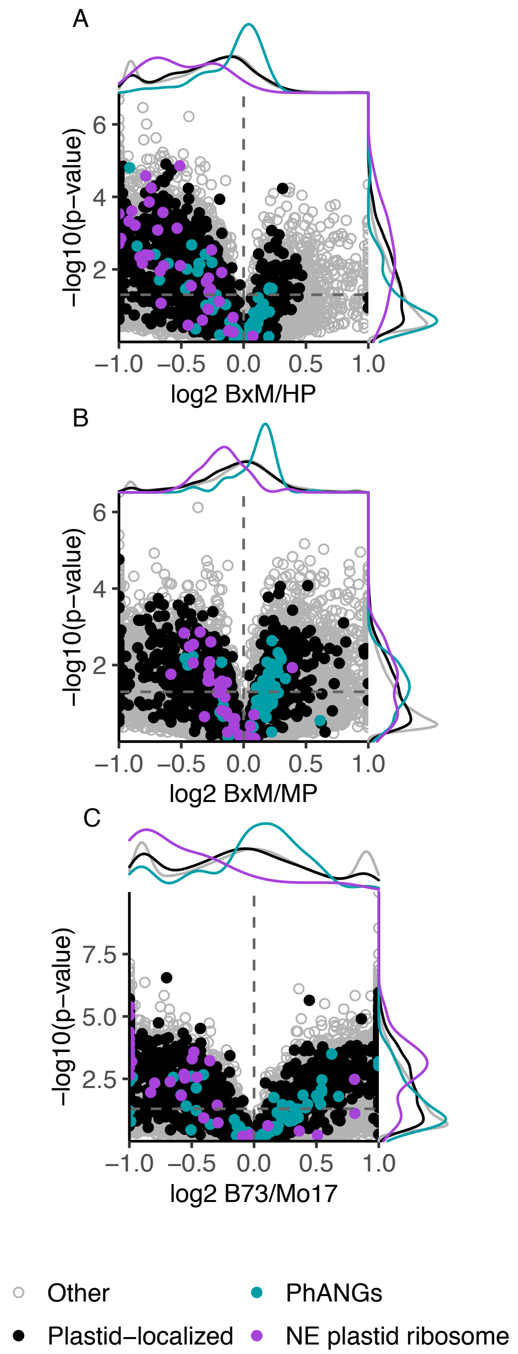


**Fig. S3.** Volcano plots displaying RNA expression patterns of the most significantly non-additive proteins, representing B73xMo17/high-parent (A), B73xMo17 /mid-parent (B), and B73/Mo17 (C). Photosynthesis-associated Nuclear Genes (PhANGs), Nuclear-Encoded (NE) plastid ribosome, and are color-coded.

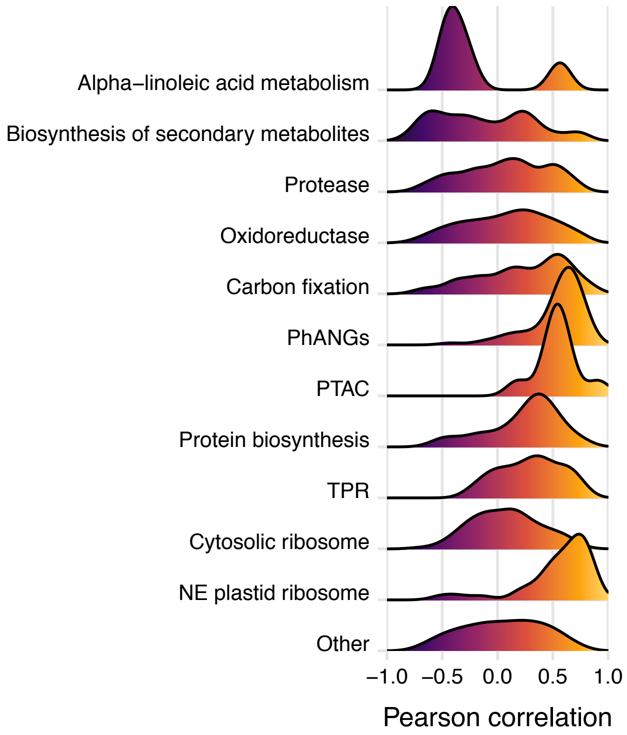


**Fig. S4.** Density curves of Pearson correlations between RNA expression heterosis and plant height heterosis in the combined RIL and six hybrids datasets.

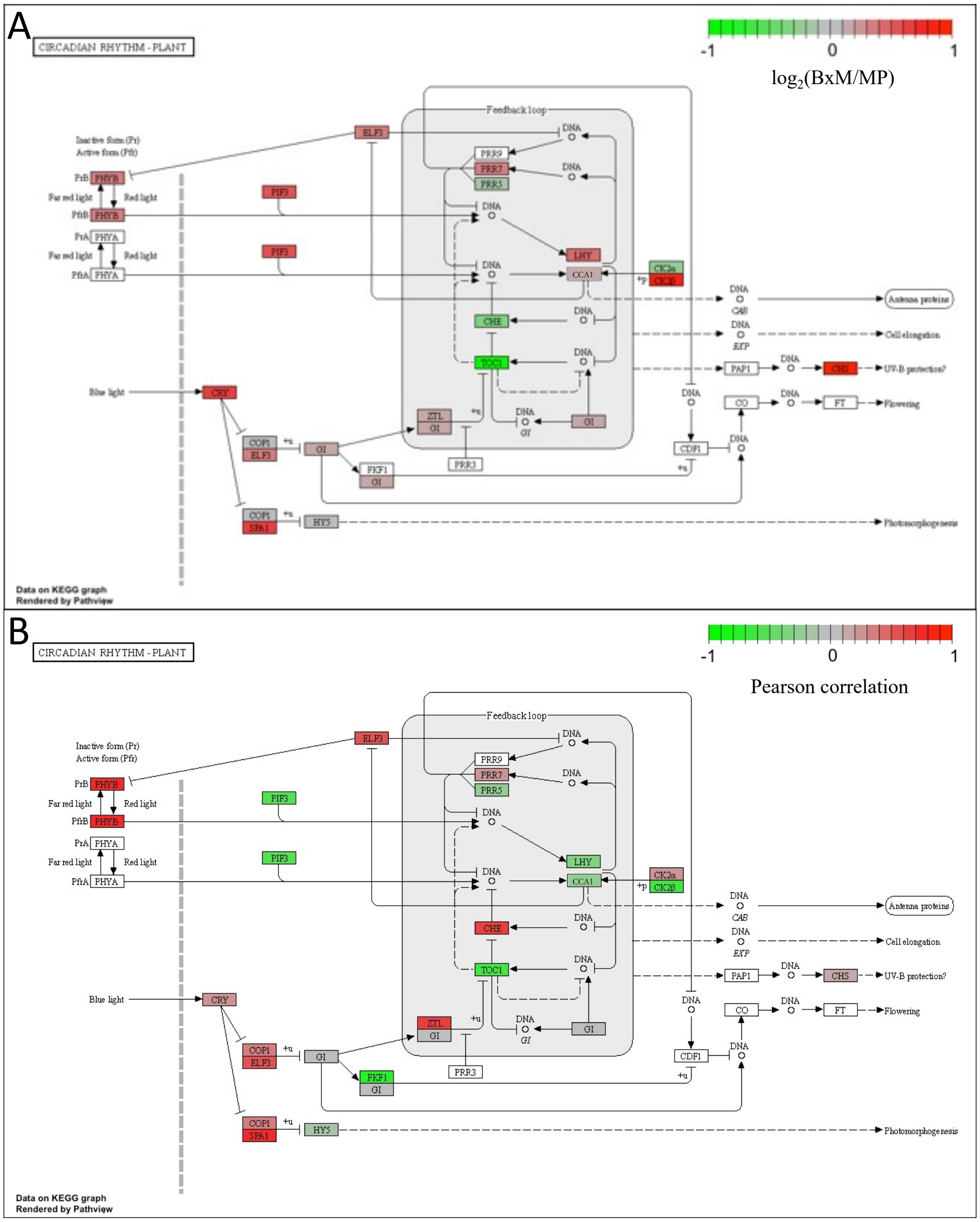


**Fig. S5.** (A) Expression of circadian rhythm-related transcripts in the hybrid, B73xMo17, relative to its mid-parent level. (B) Pearson correlation of circadian-related transcripts between expression heterosis and plant height heterosis.

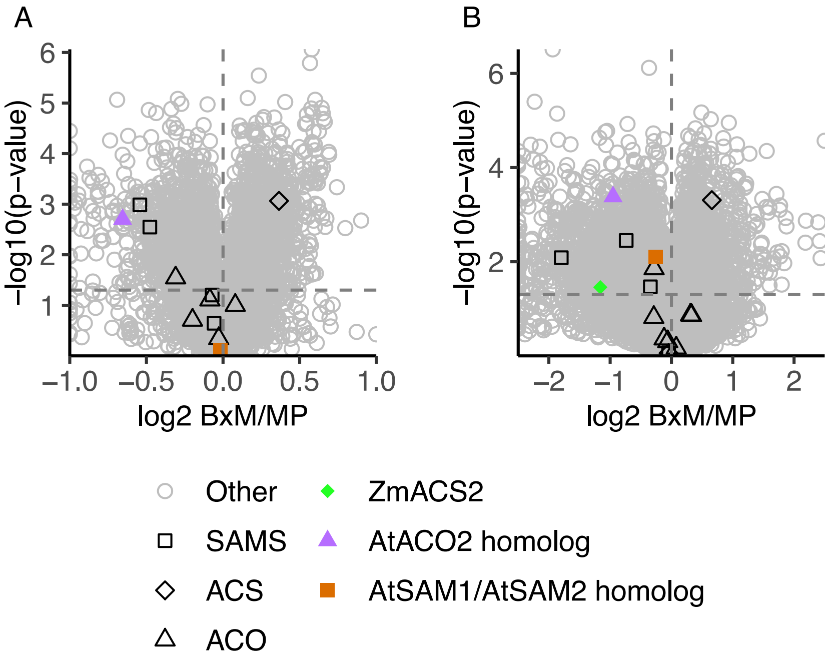


**Fig. S6.** Volcano plots showing expression heterosis of ethylene biosynthesis enzymes in the proteome (A) and transcriptome (B) of the BxM hybrid. Of the six enzymes for which the mutant is known to produce reduced ethylene (ZmACS2, ZmACS6, AtACO1, AtACO2, AtSAMS1, and AtSAMS2), two were detected in the proteome (the AtACO2 homolog and the AtSAMS1/AtSAMS2 homolog) and three were detected in the transcriptome (ZmACS2, the AtACO2 homolog and the AtSAMS1/AtSAMS2 homolog).

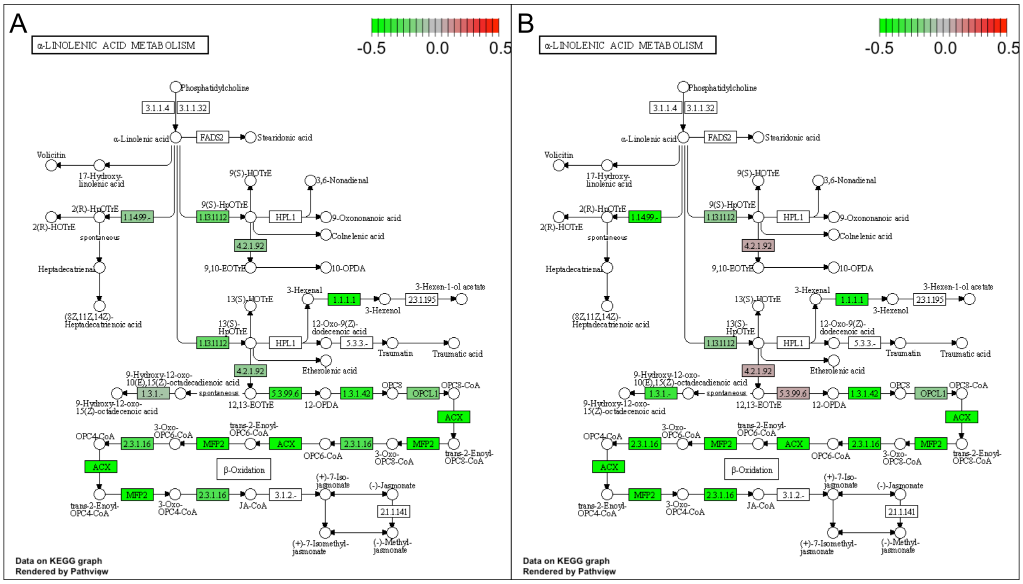


**Fig. S7.** Expression of JA biosynthesis enzymes in the (A) hybrid B73xMo17 relative to its mid-parent level; and (B) in the ET biosynthesis mutant (acs2-6) compared to its inbred B73 background.

**Dataset S1 (separate file).** Hybrid/mid-parent and parent/parent ratios for protein expression from the B73xMo17 hybrid and its parents.

**Dataset S2 (separate file).** Chromosomal breakpoints of the RILs and their hybrids. a=B73, b=Mo17, h=heterozygous.

**Dataset S3 (separate file).** Average plant heights of the RILs, diverse inbreds, and each of their hybrids used for correlation analyses.

**Dataset S4 (separate file).** Pearson correlations between expression heterosis (hybrid/mid-parent) of each protein and transcript and plant height heterosis (hybrid/mid-parent plant height).

**Dataset S5 (separate file).** Normalized tandem-mass tag (TMT) values for all experiments in this study.

**Dataset S6 (separate file).** Counts per million (CPM) values from RNA-seq analysis of seedling leaf tissue for three biological replicates each of six maize hybrids and their inbred parents.

**Dataset S7 (separate file).** Counts per million (CPM) values from RNA-seq analysis of seedling leaf tissue for three to four biological replicates each of eight maize RIL hybrids and their RIL parents, as well as the B73xMo17 hybrid and its parents.

**Dataset S8 (separate file).** Protein group assignments and their sources.
